# *Mycobacterium intracellulare* ABSURDO is a novel clinical isolate with three colony morphotypes that vary in pathogenicity and sequence at the *PKS* and *MtrA* loci

**DOI:** 10.1101/2025.02.03.636325

**Authors:** Tiffany A. Claeys, Yijing Liu, Anneka Prigodich, Emily Peck, Emily Gokul, Nine Reed-Mera, Albright Tuah, Abena Wirekoh, Kylie Quinn, Oscar Rosas Mejia, Gillian Clary, Smitha Sasindran, Sheri Dellos-Nolan, Erin Gloag, Anna R. Huppler, Natalie M. Hull, Brooke A. Jude, Richard T. Robinson

## Abstract

*Mycobacterium intracellulare* is a nontuberculous mycobacteria (NTM) species which can cause serious and sometimes fatal disease in immunocompromised individuals. Other NTM species, including *M. avium* and *M. abscessus*, commonly exhibit two colony morphotypes (smooth and rough) which vary in appearance and liquid growth properties. Here we characterize a novel clinical isolate of *M. intracellulare* which exhibits three (not two) colony morphotypes which differ in appearance, liquid growth properties, acid-fastness and *in vivo* survival following infection of mice via an inhalational exposure model. The genome of this isolate, which we have termed ABSURDO, as well as the genome of each morphotype components, aligns with that of *M. intracellulare* yet contains ∼16% more protein coding sequences than the *M. intracellulare* type strain ATCC 13590^T^. Variation analysis of each morphotype genome revealed that across the three morphotypes there were only two mutations which had a high likelihood of causing a phenotype due to a genetic change: one in the gene encoding modular polyketide synthase (PKS), and another in the two component system response regulator MtrA. Neither of these genes have been previously implicated in the morphotype shifting of an NTM. In summary, *M. intracellulare* ABSURDO is a novel pathogenic isolate with a genome that aligns with (but is nevertheless larger than) the *M. intracellulare* type strain and comprises three morphotype components which differ in two genes that have not been implicated in NTM appearance, acid-fastness, *in vitro* and *in vivo* growth.

## INTRODUCTION

Nontuberculous mycobacteria (NTM) represent >170 species of the genus *Mycobacterium* (1). NTM are ubiquitous in natural environments such as soil and water, as well as anthropogenic environments such as water mains, shower heads, swimming pools, and household dust (2, 3). Although they share the genus with the more notorious pathogen *M. tuberculosis*, NTM have until recently been comparatively neglected as a focus of basic research. Increasing rates of NTM infection in many regions have, however, raised popular awareness of NTM (4, 5) and elevated the significance of researching these opportunistic pathogens and developing new treatment options (6, 7). Clinical manifestations of NTM infection include pulmonary disease as well as skin and soft tissue, lymph node, musculoskeletal, or disseminated infections (8). The reasons why NTM disease incidence and/or prevalence are increasing are unknown, but may be tied to climate change as reports of NTM disease are higher among tropical regions (9–11), the range of which is widening (12, 13), and regional NTM incidence can increase following extreme weather events and environmental disruptions (14–18).

Three NTM species that commonly cause human disease are *M. abscessus*, *M. avium*, and *M. intracellular*e. In clinical microbiology terms, *M. abscessus* is a rapid-growing mycobacteria that typically appears within 2-3 days of culture on agar media. *M. avium* and *M. intracellulare*, on the other hand, are slow-growing mycobacteria that appear within 3-4 weeks of culture on agar media. *M. avium* and *M. intracellular*e are not often speciated or otherwise distinguished from one another in a clinical microbiology lab setting; instead, they are often collectively referred to as either *M. avium* complex (MAC) or *M. avium-intracellulare* (MAI). This lack of distinction is unfortunate, however, as *M. avium* and *M. intracellulare* differ in their infectivity characteristics, virulence, clinical features, and antibiotic sensitivities. Although *M. avium* has historically been the more studied species of the two, *M. intracellulare* is estimated to account for 45-50% of all MAC lung disease diagnoses (19, 20). The two species have significant differences in drug susceptibility, with *M. intracellulare* displaying a lower resistance to clinically relevant antibiotics than *M. avium* (21, 22). *M. avium* is the species most associated with a concurrent AIDS diagnosis, accounting for over 95% of MAC infections in AIDS patients compared to only ∼50% in non-AIDS patients (23). Clinically, patients infected with *M. intracellulare* exhibit a more severe and advanced form of disease than those infected with *M. avium*, more often presenting with the more severe fibrocavitary form of disease, having a lower treatment response rate, and generally having a worse overall prognosis (19). Among individuals with cystic fibrosis, the hospital acquired transmission characteristics of *M. avium* versus *M. intracellulare* also differ (24). It is therefore important to understand the underlying pathogenesis and nuances of the host immune response specific to *M. intracellulare*.

A peculiar feature of numerous NTM isolates is their appearance on solid growth media as colonies with more than one morphology, or morphotype (25). The two most common colony morphotypes are described as “smooth” versus “rough” (26). The smooth morphotype is characterized by a uniform and glossy appearance, in contrast to rough morphotype colonies which appear irregular, dry, and corded. The occurrence of smooth and/or rough colonies distinguishes NTM isolates from *M. tuberculosis* isolates, which are uniformly rough. The biochemical basis for a smooth versus rough morphotype are differences in membrane lipid composition (27–29), and depending on the species, smooth and rough NTM morphotypes can exhibit virulence differences in animal models (30–33). In *M. abscessus*, mutations in the genes that encode mycobacterial membrane protein large 4 (*Mmpl4*) and the gene cluster *mps1-mps2-gap* underlie smooth-to-rough morphotype shift of *M. abscessus* (34, 35). *Mmpl4* and glycopeptidolipid-addressing protein (*GAP*) aid the transport of glycopeptidolipids (GPLs) across the cell membrane (36, 37), and Mps1 and Mps2 promote the synthesis of GPLs (38). therefore, mutations in the genes encoding these proteins can lead to differences in lipid export efficiency and metabolism, which in turn influence the cell wall composition and overall virulence of *M. abscessus*. Whether mutations in the same genes are responsible for the morphotype variation in other NTM species such as *M. intracellulare*, or instead are *M. abscessus*-specific, is unknown as no such whole genome comparisons of *M. intracellulare* morphotypes have been made.

Here we characterize a novel clinical isolate of *M. intracellulare* that exhibits three (not two) distinct colony morphotypes. As we show, each morphotype differs in macroscopic and microscopic appearance, acid-fastness, liquid growth properties and *in vivo* survival upon infecting mice via inhalation exposure. Whole genome sequencing of the isolate and each morphotype component demonstrated this isolate, which we have named ABSURDO, to be a single species (not a mixture) with morphotypes distinguished by mutations in the genes modular polyketide synthase (PKS) and the two-component response regulator MtrA, thus distinguishing *M. intracellulare* morphotypes from *M. abscessus* morphotypes in genetic etiology.

## METHODS

### Ethics statement

All animal studies were reviewed and approved by The Ohio State University (OSU) Institutional Animal Care and Use Committee (IACUC). All experiments involving *M. intracellulare* followed procedures and protocols that were approved by the local Institutional Biosafety Committees (IBC) at Medical College of Wisconsin, OSU and BARD College.

### Mice

C57BL/6 and NFκB-luciferase reporter mice (FVB.Cg-Tg(HIV-EGFP,luc)8Tsb/J) (39) were purchased from Jackson Laboratory (Bar Harbor, ME) and housed within the AALAC-accredited University Laboratory Animal Resources facility.

### Bacterial culture

*M. intracellulare* ABSURDO (an anagram of OSU and BARD) is a clinical isolate that originated from the cervical lymph node of a child being treated for NTM lymphadenitis (40). Our immune profiling of other children with NTM lymphadenitis has been prevouisly reported (40).Originally collected on Lowenstein–Jensen agar, for our study the isolate was maintained and cultured on Middlebrook 7H10 agar supplemented with oleic acid dextrose complex (OADC), as produced per the protocols of Ordway and Orme (41). Consistent with the classification of *M. intracellulare* as slow growing mycobacteria, colonies of *M. intracellulare* ABSURDO and its morphotype components only appeared after 3-4 weeks of culture (37°C, 5% CO_2_). As described in our *Results*, three distinct colony morphotypes were distinguishable upon culture of the original isolate, which we also refer to as the Parent (P). To generate pure cultures of each morphotype, colonies of each were subcultured via streaking onto fresh plates and incubating for an additional 3-4 weeks. This process was repeated several times over the course of 2-3 months until we attained plates with uniform colony appearance. We refer to the three morphotypes as Component A, Component B, and Component C. *M. intracellulare* ABSURDO and its three morphotypes were initially speciated by the OSU Wexner Medical Center Clinical Microbiology Laboratory via matrix-assisted laser desorption/ionization mass spectrometry (MALDI-MS). Glycerol stocks of P and Components A-C were stored at −80°C; subsequent thawing of the frozen stocks and re-plating showed that all remained viable, retained pure colonies, and showed no evidence of morphotype-switching.

### Acid fast staining, visualization, and quantitation

Slides with smears of *M. intracellulare* ABSURDO and its morphotype components were prepared and acid fast stained per established protocols (42). To quantify the percent of bacilli which were acid fast positive (AF^POS^) versus acid fast negative (AF^NEG^), we first took digital images of each slide and morphotype at 100× magnification with oil immersion. We then drew identically sized squares around seven regions of interest (ROIs), a ROI being defined as ∼50-100 bacilli that were sufficiently separated from one another (i.e. not clumped together) to enable distinction of whether an individual bacillus was red (AF^POS^) or blue (AF^NEG^). The percent AF^POS^ per ROI was then calculated.

### DNA extraction and sequencing

*M. intracellulare* ABSURDO and its morphotype components were individually cultured in 7H9 media supplemented with OADC for ∼7 days (37°C, shaking at 220 RPM), after which DNA was extracted per previously described methods (43) and then shipped to a commercial sequencing service (SeqCoast Genomics, Portsmouth, NH). There, DNA samples were prepared for whole genome sequencing using an Illumina DNA Prep tagmentation kit and unique dual indexes. Sequencing was performed on the Illumina NextSeq2000 platform using a 300-cycle flow cell kit to produce 2×150bp paired reads. 1-2% PhiX control was spiked into the run to support optimal base calling. Read demultiplexing, read trimming, and run analytics were performed using DRAGEN v3.10.12, an on-board analysis software on the NextSeq2000.

### Sequence assembly and annotation

Sequence files were uploaded to Galaxy EU platform for further analysis. Files were trimmed using FastP (Galaxy Version 0.23.2+galaxy0) and the standard parameters. Resulting trimmed files were assembled using Unicycler, removing contigs smaller than 500 bp in length (Galaxy Version 0.5.0+galaxy1). Assembly quality was assessed via Quast (Galaxy Version 5.2.0+galaxy1). Contigs were annotated using Prokka (Galaxy Version 1.14.6+galaxy1). Presence of genes that produce secondary metabolites were screened using Antismash (Galaxy Version 6.1.1+galaxy1). Contigs were downloaded and uploaded to BV-BRC (v. 30.13.9) for further characterization using the comprehensive genome analysis service (https://www.bv-brc.org).

### Whole Genome Sequence Accession

The whole genome shotgun (WGS) sequencing projects described in this report have been deposited at DDBJ/ENA/GenBank under the following accession numbers: JAWLLF000000000 (this WGS corresponds to the Parental isolate), JAWLLE000000000 (corresponds to Component A), JAWLLD000000000 (corresponds to Component B), and JAWLLC000000000 (corresponds to Component C).

### Phylogenetic Analysis

Phylogenetic analysis was performed by the BV-BRC genome analysis service. The closest reference and representative genomes to were identified by Mash/MinHash (44). PATRIC global protein families (PGFams) (45) were selected from these genomes to determine the phylogenetic placement of this genome. The protein sequences from these families were aligned with MUSCLE (46), and the nucleotides for each of those sequences were mapped to the protein alignment. The joint set of amino acid and nucleotide alignments were concatenated into a data matrix, and RaxML (47) was used to analyze this matrix, with fast bootstrapping (48) was used to generate the support values in the resulting phylogenetic tree.

### Bone marrow derived macrophage culture

To generate bone marrow derived macrophages (BMDMs), bone marrow cells were collected from the femurs and tibias of NFκB-GFP-luciferase reporter mice (FVB.Cg-Tg(HIV-EGFP,luc)8Tsb/J) and cultured as previously described (49). For each experiment, BMDMs were seeded at 5 × 10^5^ cells per well in a 6 well plate and allowed to adhere for 18 hours before stimulation with P or Components A, B, or C.

### *M. intracellulare* morphotype BMDM stimulation assay

Glycerol stocks of P and Components A, B and C were thawed, diluted, and counted using a Petroff-Hausser chamber. Since mycobacteria tend to clump, all bacteria (P, A, B, and C) were counted after first allowing clumps to precipitate out of solution using a ‘third tube method’ established by Hall-Stoodley et al (50). P, A, B, and C were then individually added to the wells containing BMDMs (MOI: 10) and incubated in a humidified 5% CO2 incubator at 37°C. After 48 hours, the supernatants were removed (for ELISA analysis of TNFα levels) and the BMDMs were lysed for luciferase activity measurements using the Dual-Luciferase Reporter Assay System (Promega, Madison, WI). TNFα levels were measured using the Biolegend ELISA MAX Deluxe method (San Diego, CA).

### *M. intracellulare* infection via inhalation exposure

Four groups of C57BL/6 mice were infected with each individual *M. intracellulare* preparation (P, A, B, or C) via inhalation exposure, using the Schuco Compressor Nebulizer system (Model S5000). Inside a biosafety cabinet, an inoculum of 10^8^ CFU (i.e. 10 mL of a 10^7^ CFU/mL solution in sterile water) was pipetted into the nebulizer portion and then atomized into a chamber containing the mice (the chamber also being inside the biosafety cabinet). This was done simultaneously for each morphotype, with one morphotype per aerosol system to prevent cross contamination, so we could compare their *in vivo* growth kinetics at the same time and using the same batches of 7H10/OADC plates and other reagents (to eliminate batch effects). Each group of mice had roughly equal portions of male and females. On the indicated days post-infection, animals were euthanized, and their lungs were removed, homogenized in normal saline using the gentleMACs tissue dissociation system (Miltenyi Biotec), and plated on 7H10/OADC plates. Plates were incubated in a humidified 5% CO2 incubator at 37° for 3-4 weeks before counting CFUs. In addition to determining CFU burden at each time point, we also documented the proportion of colonies that exhibited each morphotype.

### Statistics and graphs

Graphs and associated statistical analyses were prepared using GraphPad Prism (version 9.0) with One Way ANOVA and Tukey’s multiple comparisons test (for normally distributed data), Brown-Forsythe and Welch ANOVA with Dunnett T3 multiple comparison test (for data that were not normally distributed), or Student’s t-tests for less than three groups. Differences were considered significant if p < 0.05 and are graphically represented by an asterisk.

## RESULTS

### I. The isolate *M. intracellulare* ABSURDO comprises at least three morphotypes which vary in colony appearance and liquid growth properties

The isolate *M. intracellulare* ABSURDO was originally cultured from the cervical lymph node of a child who had been treated for NTM lymphadenitis (40). For the duration of our study the isolate and its morphotype derivatives were maintained on 7H10 supplemented with OADC. The isolate had an off-white appearance that became yellow with age (**FIG 1G**) and was positively identified via clinical mass spec analysis as *M. intracellulare* (MALDI Score=2.21). Upon close visual inspection of the parent (P) isolate, we noted three distinct colony morphologies that we streaked to isolation and designated Components A, B and C (**FIG 1A-C**). Component A exhibited features of the canonical “smooth morphotype” observed in other NTM species (51–53): round shape, smooth margins, a convex (a slight dome) elevation, white with a glistening appearance (**FIG 1A**). Low-magnification examination of the Component A margins showed translucent, branched matrices around the entire colony perimeter (**FIG 1D**). Component B exhibited classical “rough morphotype” characteristics: a larger colony with undefined edges and a dull appearance (**FIG 1B**). Component C exhibits an intermediate phenotype between the smooth and rough morphotypes: a domed, glistening appearance, but with scalloped edging more commonly seen with rough morphotypes (**FIG 1C**). Unlike Component A, neither Component B nor Component C had branched matrices along their perimeters (**FIG 1E-F**). Like the original isolate P (**FIG 1G**), our streak-purified cultures of Component A, B, and C were also identified as *M. intracellulare* by mass spec analysis (MALDI scores=2.12[A], 2.08[B], and 2.00[C], respectively). Serial dilutions of our P glycerol stock were used to estimate the relative abundance of each morphotype as follows: ∼50% Component A; ∼35% Component B; ∼15% Component C.

**FIGURE 1.**
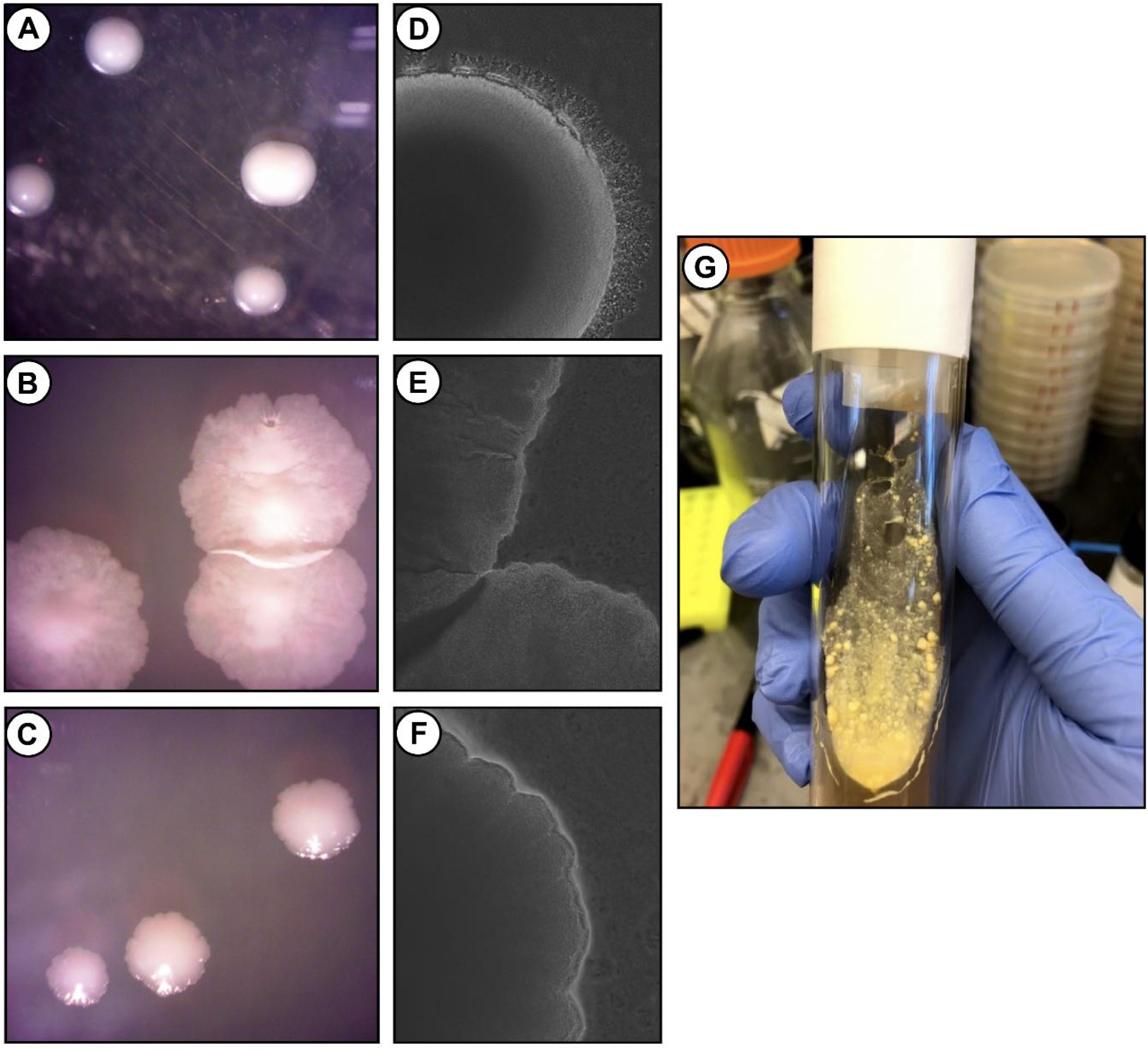
*M. intracellulare* ABSURDO comprises three morphotypes which vary in colony appearance. The clinical isolate *M. intracellulare* ABSURDO comprises three colony morphotypes which were picked and plated separately until purified stocks were obtained. (**A-C**) Representative images of the (**A**) Component A, (**B**) Component B, and (**C**) Component C morphotypes as they appear on 7H10/OADC. (**D-F**) Low magnification stereomicroscope images of the colony borders of (**D**) Component A, (**E**) Component B, and (**F**) Component C. (**G**) The original or parent (P) isolate of *M. intracellulare* ABSURDO.

*M. intracellulare* ABSURDO morphotypes were also distinct from one another with regards to their liquid growth properties (**FIG 2**). Shown are the results of two independent biological replicates (representative of four independent biological replicates), wherein the original isolate P and each morphotype were added to liquid media (7H9+OADC) and followed longitudinally, with growth monitored using both (OD_600_) and colony forming unit (CFU) concentration as readouts. After inoculating liquid media with P, the OD_600_ did not raise above control (i.e. no bacteria) levels until at least 73 hr post-inoculation; after this point the OD_600_ continued to increase slowly until the final time point at 217.5 hrs. The CFU concentration of P likewise increased during this period (**FIG 2B**) and generally tracked well with OD_600_ (**FIG 2C**). The liquid growth properties of A were similar to P, as measured by both OD_600_ and CFU (**FIG2A-B**), and the correlation between OD_600_ and CFU was likewise generally positive (**FIG 2C**). Contrasting with Component A were the properties of Component B, as the OD_600_ curves of Component B lagged behind that of Component A in both experimental replicates, and did not increase over control levels until 101 hours and by the final time point had only reached ∼50% the OD_600_ levels of A. The CFU curves of B lagged behind that of A for Replicate 1, but not Replicate 2 (**FIG 2B**). Consequently, the correlation between OD_600_ and CFU was not strong (**FIG 2C**), possibly due to differences in optical properties of the two morphotypes. Neither OD_600_ nor CFU concentration data are provided for Component C as liquid growth results were inconsistent across labs, whereas the growth patterns of P, A and B were consistent across three labs (NH, RR, BJ); generally speaking, Component C either failed to grow or grew at an even slower rate than Component B (data not shown). To conclude, the liquid growth rates *M. intracellulare* ABSURDO and its morphotypes can be summarized as P ≅ A > B > C.

**FIGURE 2.**
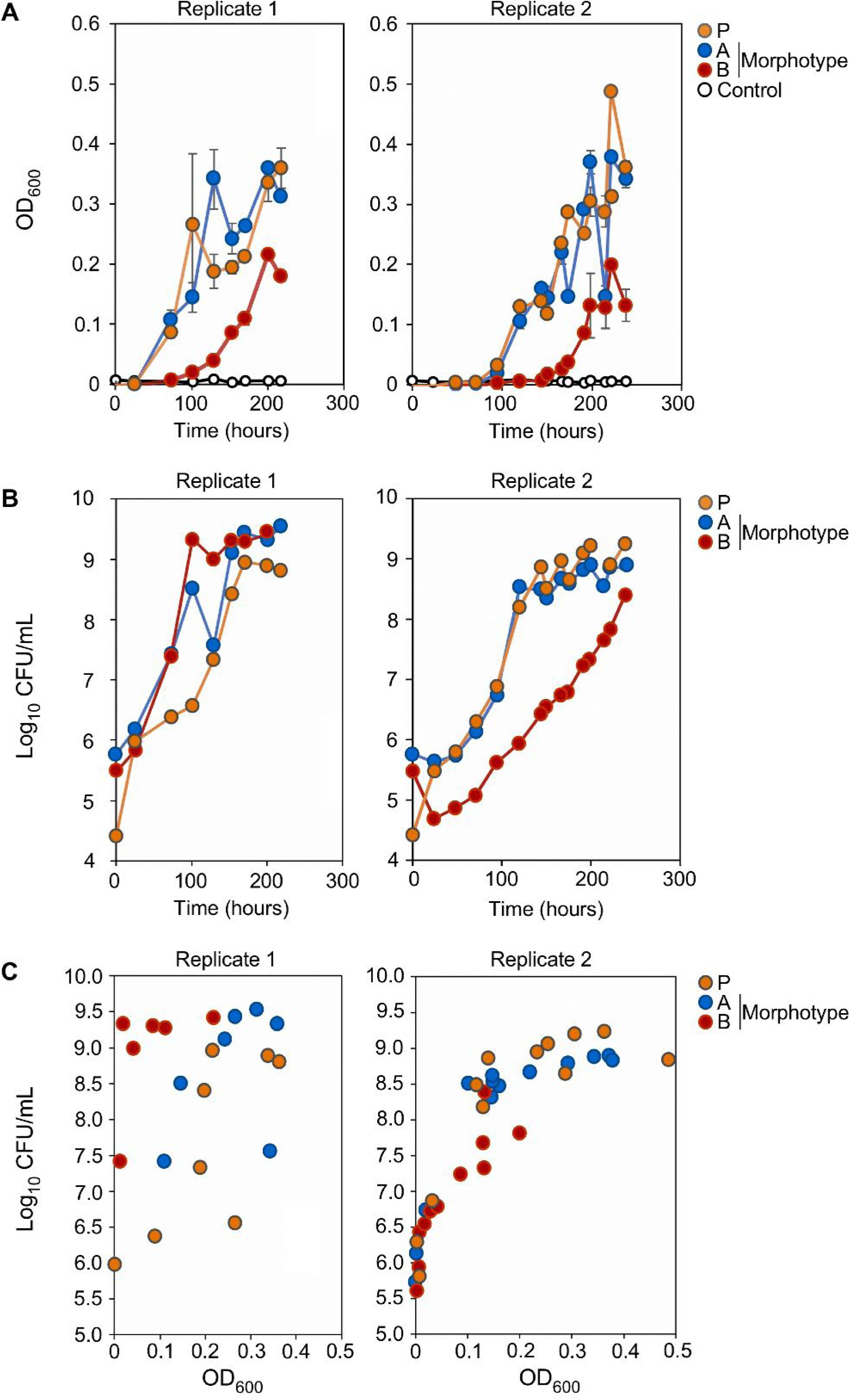
The three morphotypes of *M. intracellulare* ABSURDO vary in liquid growth properties. The original parent (P) isolate of *M. intracellulare* ABSURDO and each of its three morphotypes (Components A, B, and C) were inoculated into fresh 7H9/OADC media and cultured for 9 days. The growth of each culture was periodically measured using two readouts: (**A**) CFU concentration and (**B**) OD_600_. CFU concentration was determined by plating serial dilutions of the culture from the indicated timepoints on 7H10 plates; OD_600_ was measured using a spectrophotometer. (**C**) The relationship between CFU concentration and OD_600_ for each culture. This experiment was repeated four times. Shown are data from two independent replicates, Replicate 1 and Replicate 2.

### II. Each *M. intracellulare* ABSURDO morphotype has distinct acid staining properties and shapes

To assess their acid-fast staining profile and microscopic appearance, colonies of Components A-C were streaked onto glass slides, stained and counterstained to visualize acid-fast positive (AF^POS^) and acid-fast negative (AF^NEG^) bacilli. Slides of each morphotype were stained at the same time to avoid batch effects and enable direct comparison of staining patterns. Digital images of each slide were used to quantify the percent AF^POS^ bacilli within several fields of view (see *Methods*). Among the three morphotypes, Component A had the highest and most consistent degree of acid-fast staining (**FIG 3A, E**). Most Component B bacilli were also acid fast (**FIG 3B**); however, AF^NEG^ bacilli were also evident, resulting in a significant decline in percent AF^POS^ values (**FIG 3E**). The extent of acid fastidiousness was even lower for Component C (**FIG 3C**), which had the lowest percent AF^POS^ bacilli of all morphotypes (**FIG 3E**). Another noteworthy feature of Component C that distinguished it from other components: the individual bacilli of Component C were more likely to appear as part of a chain of 4-14 individuals, akin to *Streptococci* species (**FIG 3C**). Consistent with P consisting mostly of A and B, most P bacilli were AF^POS^ (**FIG 3D, E**). The above properties were not unique to the colony form of each component, as similar results were obtained using liquid cultures of each morphotype (**FIG 3F-I**).

**FIGURE 3.**
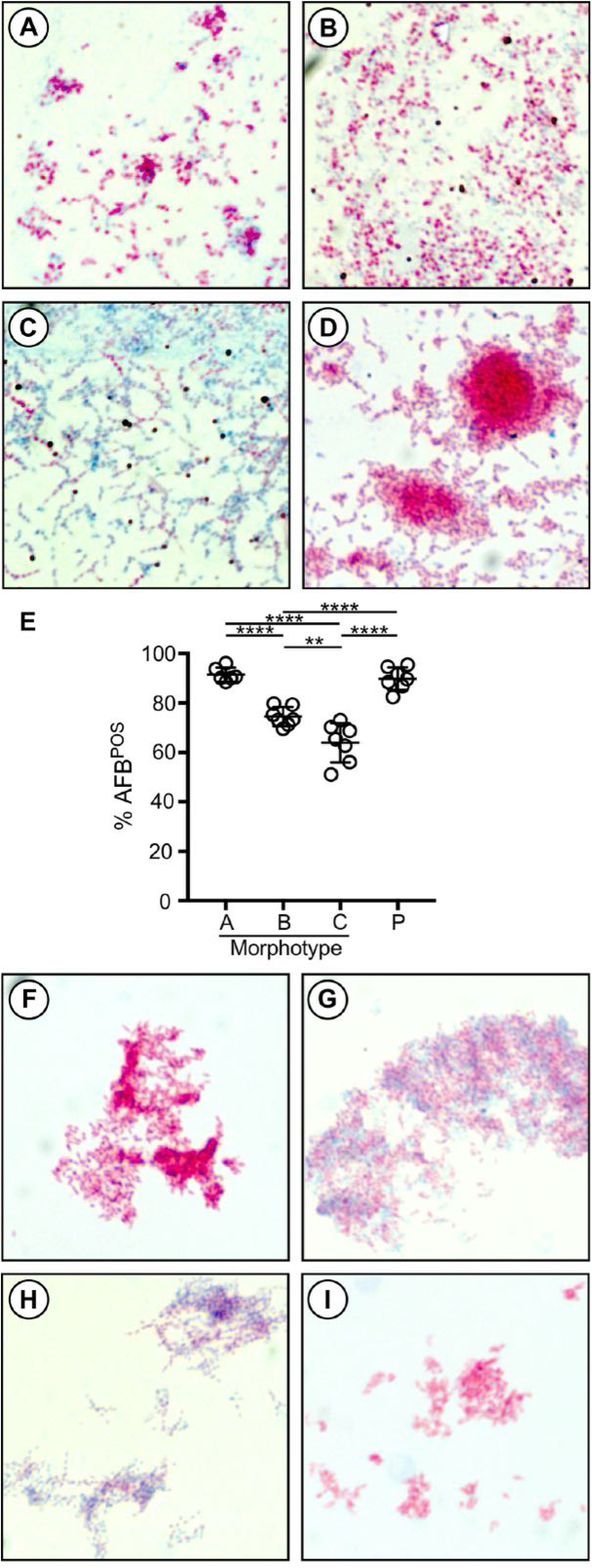
The three morphotypes of *M. intracellulare* ABSURDO vary in microscopic appearance and acid fastness. (**A-E**) Colonies of each morphotype on solid agar were streaked onto glass slides and stained to visualize acid fast (AF^POS^) bacilli. Slides were imaged using oil immersion, 63X objective. Shown are representative images of morphotype (**A**) Component A, (**B**) Component B, (**C**) Component C, and (**D**) the parent (P) isolate. (**E**) The percent of bacilli within seven separate regions of interest (ROI) that were acid fast positive (%AF^POS^). Each data point represents a single ROI for the indicated morphotype or P. (**F-I**) Liquid cultures of each morphotype were treated and stained in the same manner as above. Shown are representative images of (**F**) Component A, (**G**) Component B, (**H**) Component C, and (**I**) P. Images are representative of three separate biological replicates.

### III. Mouse lung infection properties following inhalation exposure

The literature regarding *in vivo* behavior of *M. intracellulare* is limited (most NTM disease models utilize either *M. abscessus* or *M. avium*) but nevertheless demonstrates that *M. intracellulare* is capable of establishing chronic infection in mice (54, 55) ^54, 55^ and—unlike *M. tuberculosis*, which is sensitive to the presence of activated macrophages—the *in vivo* growth of *M. intracellulare* is enhanced by presence of activated macrophages (56)^56^. To assess whether *M. intracellulare* ABSURDO and its morphotype components can activate macrophages and establish a chronic infection in mice, we cultured mouse bone-marrow derived macrophages (BMDMs) in the presence of each purified morphotype and the parent isolate (MOI: 10), as well infected mice with the same preparations via inhalation exposure. For BMDM studies, we used bone marrow from transgenic NFκB-GFP-luciferase reporter mice (39) ^39^ which express intracellular luciferase reporter protein upon NFκB activation, in addition to secreting TNFα. For inhalation exposure studies, we used C57BL/6 mice and assessed lung bacterial burdens on post-infection Days 1, 48, and 124.

As shown in **FIG 4**, BMDMs respond to each morphotype by activating NFκB (**FIG 4A**) and secreting TNFα (**FIG 4B**). Contrasting with their comparable macrophage activating abilities *in vitro* were their diverging deposition and growth kinetics in vivo (**FIG 4C**). Namely, despite our loading each culture into the aerosol apparatus at the same inoculum concentrations (see *Methods*), our Day 1 lung burden assessment showed P, A, and B deposited into mouse lungs at higher levels (∼10^3^ CFU) than C (∼10^2^ CFU). By post-infection Day 48, P and A lung burdens had risen to ∼10^6^ CFU and ∼10^4^ CFU, respectively, whereas B had declined, and C was no longer detectable (**FIG 4C**). On post-infection Day 124, P was still present in the lungs at ∼10^6^ CFU; however, in mice infected with either A, B, or C alone there were no lung CFUs detected. Interestingly, whereas the lungs of mice infected P retained a mixture of morphotypes on Day 1, by Day 48 and Day 124 all the colonies were Component A (i.e. neither Component B nor Component C were noted among the lung homogenates of mice infected with P). No groups or animals showed acute signs of illness or weight loss following their initial infection (supplemental **FIG S1**). Collectively, these data demonstrate that although each morphotype can activate macrophages and infect mice on their own (*in vivo* survival of A > B > C), it is only as P that *M. intracellulare* ABSURDO survives at the highest levels and for the longest time *in vivo*, and that the P-infected lung environment selects for Component A.

**FIGURE 4.**
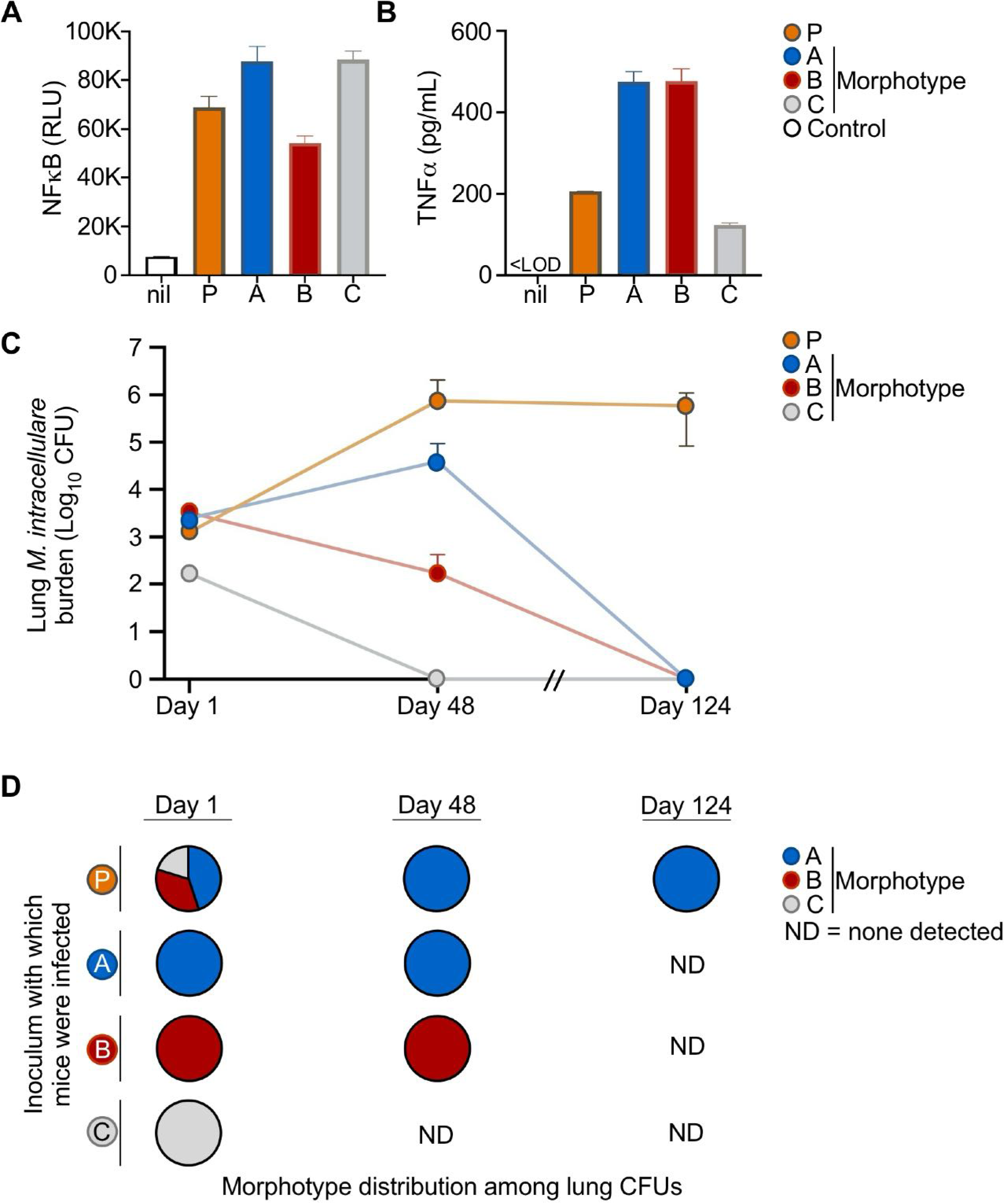
*M. intracellulare* ABSURDO morphotypes vary in their ability to establish chronic lung infection in mice. (**A-B**) Bone marrow derived macrophages (BMDMs) from NFκB-luciferase reporter mice were cultured in the presence of either media alone with no bacteria (nil), the parent (P) isolate of *M. intracellulare* ABSURDO, or its individual morphotype Components A, B, or C (MOI=10). Shown are (**A**) cellular luciferase activity and (**B**) secreted TNFα levels 48 hours later in response to each individual morphotype, with BMDMs cultured in media alone (no bacteria) serving as unstimulated controls. (**C-D**) Four groups of C57BL/6 mice were infected via inhalation exposure with either P or Components A, B, or C. On the indicated day post-infection, lungs from mice in each group (5 mice/group/timepoint) were plated on 7H10+OADC to determine (**C**) the number of lung CFUs and (**D**) within each group the proportion of CFUs which exhibited a morphotype consistent with Component A (red), Component B (blue) or Component C (gray). The results in (**A-B**) are representative of 4 independent biological replicates; the results in (**C-D**) are representative of 2 independent biological replicates.

### IV. The genome sequence of *M. intracellulare* ABSURDO

The non-uniform nature of Component B and Component C’s acid-fast staining profile (i.e. their comprising both AF^POS^ and AF^NEG^ bacilli), as well as the *Streptococci*-like appearance of Component C, raised the unsettling possibility that—despite our clinical mass spec results indicating otherwise—we were not working with a single species, but rather a mixture of two or more species. To address this, we prepared DNA from liquid cultures of P, A, B, and C for whole genome sequencing via the Illumina platform per our previously reported methods (1) ^1^. The genomes of P, A, B, and C were each sequenced via short-read sequencing and analyzed per our previous studies (57, 58). Draft assemblies of the four genomes averaged 177 contigs with L50 values ranging from 12-15, and N50 values ranging from 132,917-150,984 bp (**TABLE 1** and **FIG 5A, FIG S2A-S4A**). The genome sequences of P, A, B and C ranged from 6.06-6.08 Mbp in length (**TABLE 1**), and each most closely aligned with that of a single species: *M. intracellulare* (**FIG 6**). This genome size is slightly larger than that of *M. intracellulare* strain ATCC 13590^T^, which is 5.4 Mbp (59). Additional general features of the ABSURDO morphotypes and their comparability to ATCC 13590^T^ include similar GC content (67.7% in ABSURDO vs 68.10% in ATCC 13590^T^), a similar number of tRNA genes (48 in ABSURDO versus 47 in ATCC 13590^T^), and ∼16% more protein coding sequences or CDs (5955-5999 depending on the ABSURDO morphotype, versus 5,145 in ATCC 13590^T^) (**TABLE 1**). As with the genome of *M. avium* ss *paratuberculosis* (60), the next closest phylogenetic neighbor (**FIG 6**), most CDs in the ABSURDO genome contribute to subsystems which regulate metabolism, protein processing, stress response, defense and virulence **(FIG 5B, FIG S2B-S4B)**.

**TABLE 1.** Genome characteristics of *M. intracellulare* ABSURDO and its three morphotype components.

|  | Parent isolate | Morphotype<br>Component A | Morphotype<br>Component B | Morphotype<br>Component C |
| --- | --- | --- | --- | --- |
| **Assembly**<br>**details** |  |  |  |  |
| Contigs | 133 | 110 | 121 | 104 |
| GC content | 67.68 | 67.70 | 67.68 | 67.69 |
| Plasmids | 0 | 0 | 0 | 0 |
| Contig L50 | 15 | 13 | 15 | 12 |
| Genome length | 6,078,972 bp | 6,060,291 | 6,079,363 | 6,085,033 |
| Contig N50 | 132,917 | 148,654 | 132,917 | 150,984 |
| **Genome**<br>**features** |  |  |  |  |
| CDs | 5,993 | 5,955 | 5,993 | 5,974 |
| tRNA | 48 | 48 | 48 | 48 |
| rRNA | 3 | 3 | 3 | 3 |

**FIGURE 5.**
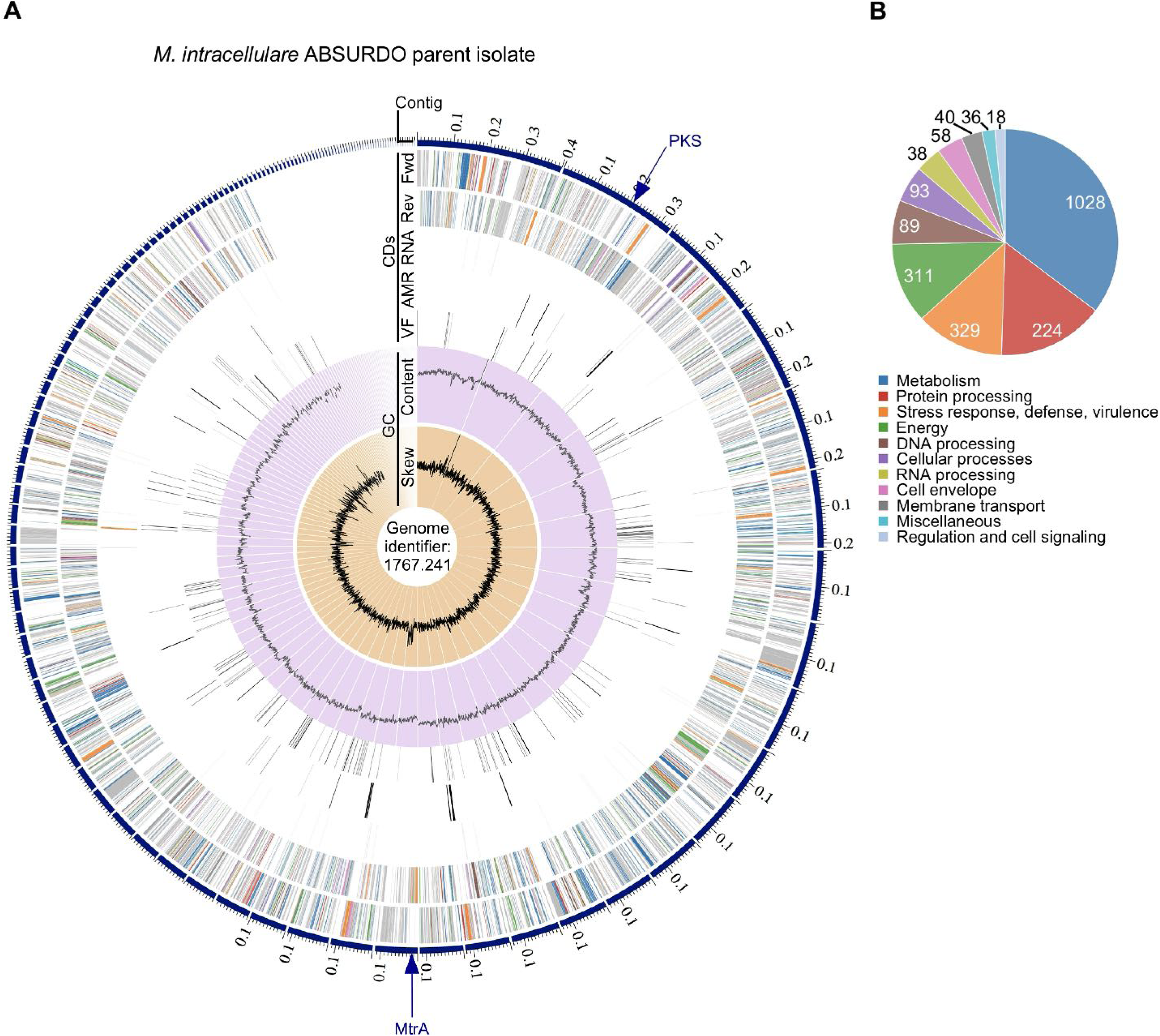
The annotated genome of *M. intracellulare* ABSURDO. Reads for *M. intracellulare* ABSURDO Parent (P) isolate were submitted to the comprehensive genome analysis service at PATRIC (81) ^82^, analyzed using RAST tool kit (82) ^83^, and assigned a unique genome identifier of 1767.241. (**A**) A circular graphical display of the distribution of the genome annotations, which includes, from outer to inner rings, the contigs, CDS on the forward strand, CDS on the reverse strand, RNA genes, CDS with homology to known antimicrobial resistance genes, CDS with homology to known virulence factors, GC content and GC skew. The colors of the CDS on the forward and reverse strand correspond to the subsystem to which each gene belongs per (**B**). Arrows indicate the locations of mutations in two genes—modular polyketide synthase PKS (location: contig 1767.241.con.0002 position 206440), and the two component system response regulator MtrA (location: contig 1767.241.con.0018 position 10400)—that are the focus of FIG 7 and FIG 8, respectively, that distinguish P from the morphotypes Component B and Component C. (**B**) An overview of the subsystems for this genome, a subsystem being a set of proteins that together implement a specific biological process or structural complex. Values indicate the number of genes in the specified subsystem.

**FIGURE 6.**
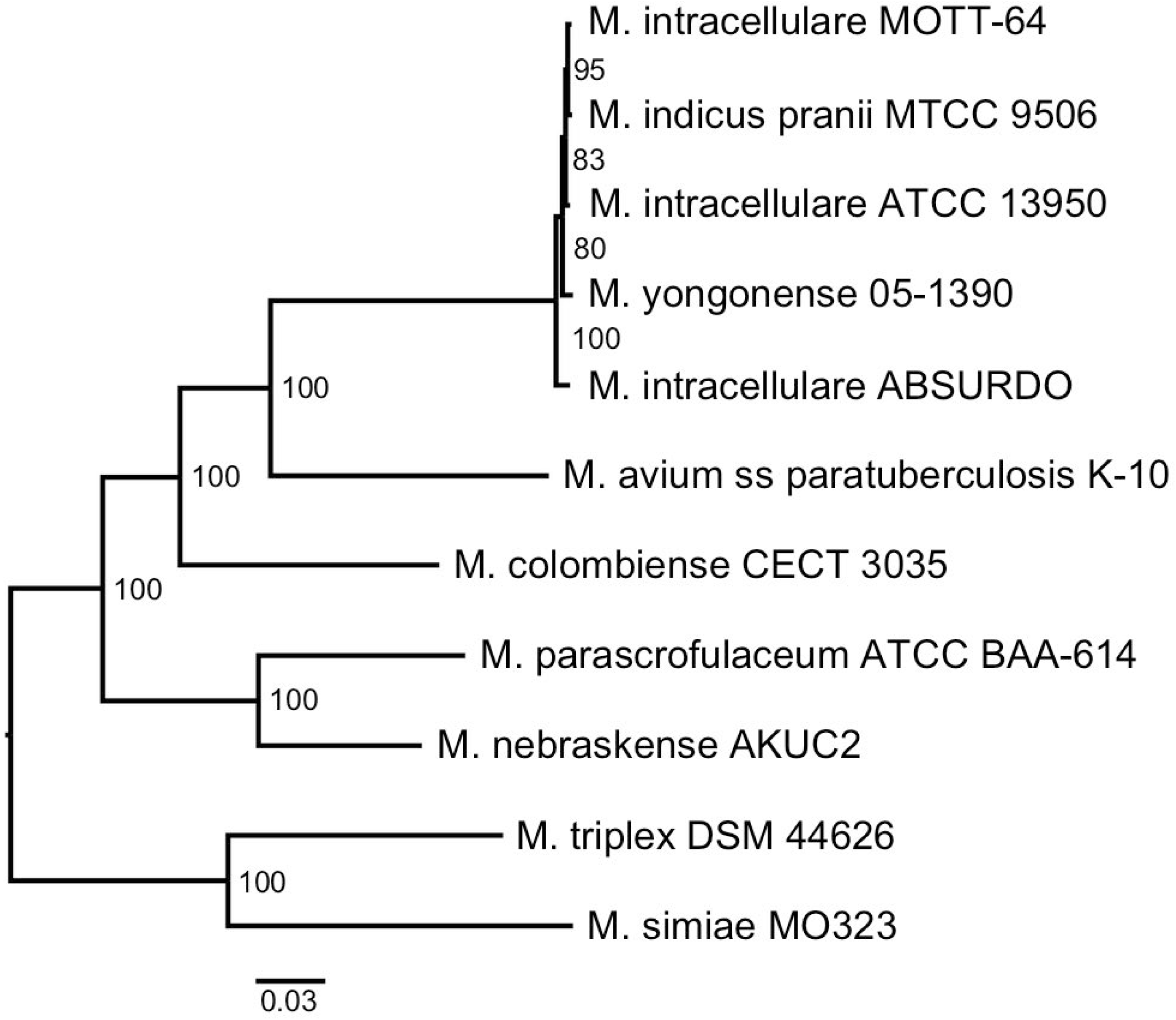
Phylogenetic analysis of the *M. intracellulare* ABSURDO genome and that of its morphotypes. Phylogenetic depiction of whole genome sequence data from the *M. intracellulare* ABSURDO parent isolate in relation to the closest reference and representative genomes. Numbers indicate the support values calculated by bootstrap analysis. Identical results were obtained when the same analysis was performed with the genome data of Component A, Component B, and Component C (data not shown).

### V. *M. intracellulare* ABSURDO morphotypes are distinguished by sequence differences in the genes PKS and MtrA

Given the differing appearances of each morphotype at the macroscopic level (**FIG 1**) and microscopic level (**FIG 3**), as well as their varying ability to grow in liquid media (**FIG 2**) and survive in lungs of infected mice (**FIG 4**), we next examined whether there were any sequence differences across morphotypes in the *MmpL* genes previously identifies as being responsible for *M. abscessus* morphotype switching (34, 35), as well as performed an unbiased variation analysis. Our findings from these genome comparisons are as follows:

- *M. intracellulare ABSURDO has few MmpL genes, none of which differ between morphotypes.* MmpLs are a subset of Resistance-Nodulation-Division (RND) transporters that translocate complex lipids and siderophores across the mycobacteria plasma membrane (61)^61^. Among MmpL substrates are cord factor (i.e. TDM, an MmpL3 substrate), long-chain triacylglycerol and mycolate wax ester (i.e. LC-TAG and MWE, both MmpL11 substrates), and mutations in *MmpL4* underlie smooth-to-rough morphotype shift of *M. abscessus* (34, 35). The number of *MmpL* genes varies depending on the species: *M. leprae* has five, *M. tuberculosis* has thirteen, and *M. abscessus* has approximately thirty putative *MmpL* genes (62). Unlike these other mycobacteria, *M. intracellulare* ABSURDO sequence had only two *MmpL* genes— *MmpL3* and *MmpL11,* which respectively were 67% and 74% identical to their *M. tuberculosis* counterparts (supplemental **SFIG S5**). There were four other CDs in the *M. intracellulare* ABSURDO genome that encode putative MmpLs and are identical in amino acid sequence to genes that were previously annotated as *M. intracellulare* RNDs (supplemental **SFIG S6**); however, since none of these were homologous to *M. tuberculosis*, *M. leprae* or *M. abscessus* MmpLs, we labeled them as putative MmpLs 1-4 (MmpL^Put1-4^, **SFIG6**). No *M. intracellulare* ABSURDO genes were identified as homologs of *M. tuberculosis MmpL4* or *M. abscessus MmpL4*, and none of the putative MmpL/RND-encoding genes were homologous to *MmpL4*; therefore, the different colony morphotypes of *M. intracellulare* ABSURDO are not due to *MmpL4* sequence differences.
- *Mutations in PKS and MtrA distinguish Component A from Component B and Component C, respectively.* Variation analysis was used to identify differences between the genomes of P, A, B and C, as well as predict the likelihood of each mutation having a low, moderate, modifier or high impact on the protein function. Among the differences observed (all are listed in supplemental **TABLE S1** and **TABLE S2**), only two genes had mutations that were predicted as having a high impact on the encoded protein function and distinguish each morphotype from one another. Relative to Component A and Component C, Component B has a frameshift mutation in modular polyketide synthase (PKS), as caused by a deletion of two nucleotides (ΔCT) from its open reading frame (**FIG 7**). Relative to Component A and Component B, Component C has a premature stop codon in the two-component system response regulator *MtrA*, as caused by a substitution (C→T) in its open reading frame (**FIG 8**). The genome of the *M. intracellulare* ABSURDO parent (P) isolate and morphotype Component A were identical at these two locations, which is consistent with Component A comprising the major morphotype of P. Potential reasons why PKS and MtrA mutations affect colony appearance and growth characteristics are included in our *Discussion*.

**FIGURE 7.**
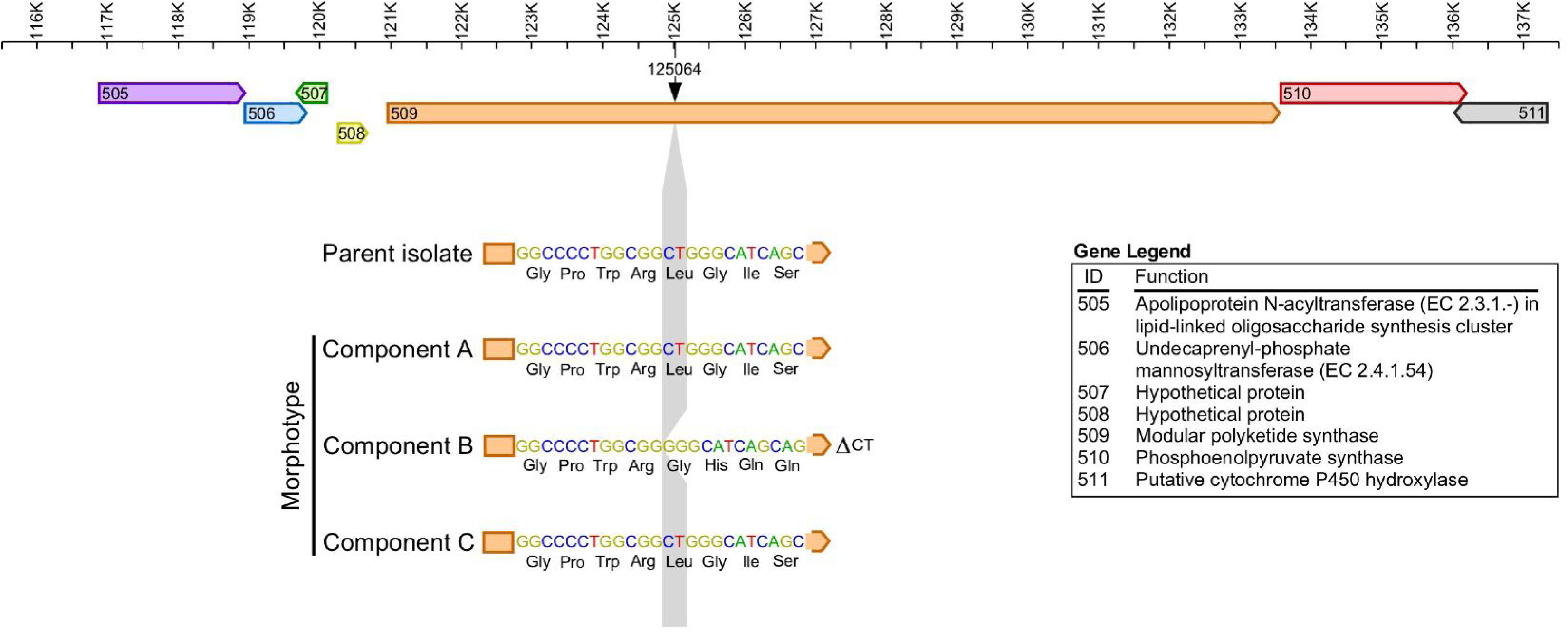
The *M. intracellulare* ABSURDO morphotype Component B has a frameshift mutation in the modular polyketide synthase (PKS) gene. A depiction of the PKS gene in relation to its closest neighbors in the *M. intracellulare* ABSURDO genome, as well as the location of a frameshift mutation (ΔCT) that distinguishes Component B from the Parent isolate, Component A, and Component C.

**FIGURE 8.**
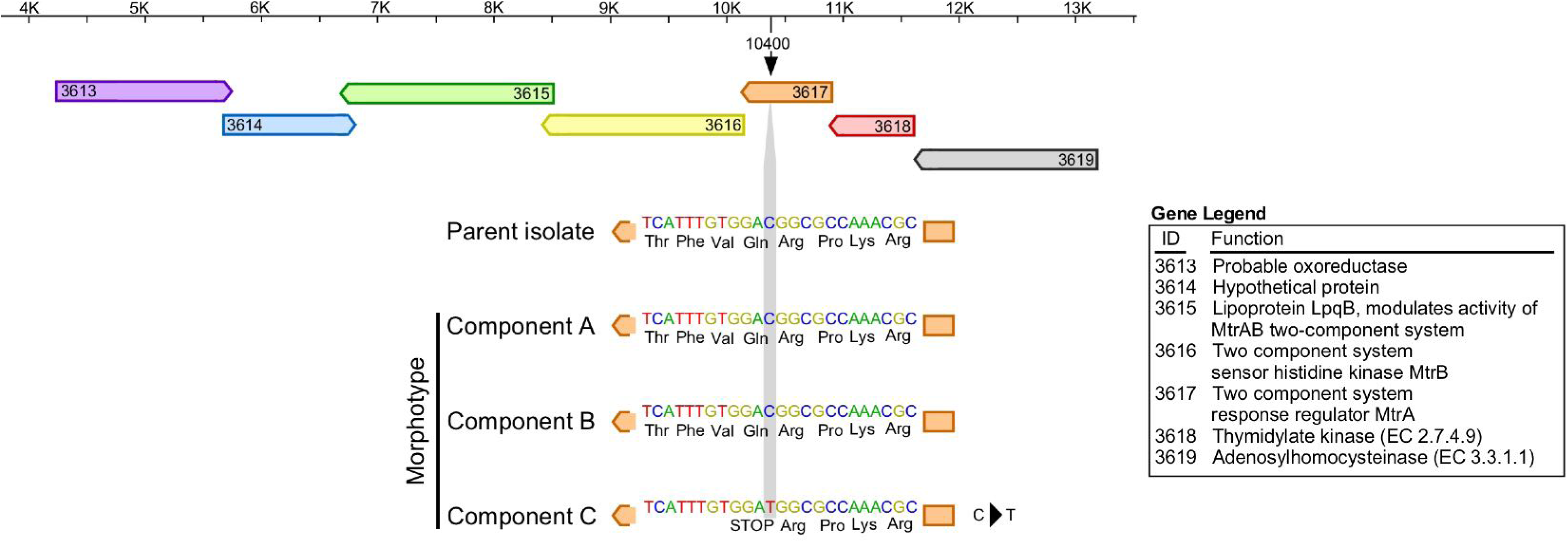
The *M. intracellulare* ABSURDO morphotype Component C has a premature stop codon in the two-component response regulator MtrA gene. A depiction of the MtrA gene in relation to its closest neighbors in the *M. intracellulare* ABSURDO genome, as well as the location of a substitution (C→T) which leads to a premature stop codon that distinguishes Component C from the Parent isolate, Component A, and Component B.

Collectively, variation analysis of each morphotype genome revealed that across the three morphotypes there were only two mutations which had a high likelihood of causing a phenotype due to a genetic change: a frameshift mutation in PKS (Component B) and a premature stop codon in MtrA (Component C). Since neither of these genes encode the MmpLs that underlie morphotype shifting *M. abscessus,* the genetic basis for *M. intracellulare* ABSURDO morphotype differences is unique from that of *M. abscessus* morphotype differences.

## DISCUSSION

*M. intracellulare* is an opportunistic pathogen that is present in the environment (63, 64) and can cause chronic infections in humans, particularly those who are immunocompromised or have an underlying lung disease such as bronchiectasis or cystic fibrosis. *M. intracellulare* can also infect otherwise healthy children and cause cervical lymphadenitis, which typically manifests as an enlarged, violaceous nodule on the neck that—although rarely painful—is disfiguring and distressing to both child and parent, and often treated with a combination of antibiotics and surgical removal (lymphadenectomy). The origin of NTM lymphadenitis is unknown but suspected to follow oral exposure to NTM during tooth eruption, enabling NTM entry into gingival tissues drained by the cervical lymph node (65). It is from cervical lymph node tissue that *M. intracellulare* ABSURDO was originally collected. *M. intracellulare* ABSURDO and its morphotypes fulfill Koch’s postulates insomuch as they were found together at a site of disease (**FIG 1G**), were isolated from the diseased tissue and grown to purity in culture (**FIG 1A-F**, **FIG 2-3**), could reestablish infection when introduced into a healthy organism (**FIG 4C**), and could be re-isolated from the experimental host (**FIG 4D**).

Although the ability of *M. intracellulare* and *M. avium* (i.e. MAC) to form different morphotypes was first reported 60+ years ago (26), the presence of a given morphotype or their ratio to one another is not currently used to inform NTM disease treatment. From a clinical perspective once a determination is made that a slow-growing mycobacteria specimen is not *M. tuberculosis*, there is (currently) minimal value ascribed to further examining it for morphotype variants such as those we describe here. We hope, however, that studies such as ours of *M. intracellulare*, as well as the extensive documentation of *M. avium* colony morphotypes by Torreles and colleagues (52, 53) and its morphotype-dependent virulence (30), will encourage closer examination of these isolates as differential morphotype representation may be predictive of whether an infection will resolve on its own in the absence of antibiotics (anti-mycobacterial drugs have numerous toxicities). Consider for example the more robust liquid growth of Component A relative to Components B and C (**FIG 2**), as well as the longer survival of Component A relative to Components B and C (**FIG 4B**); should a future clinical isolate of *M. intracellulare* comprise solely of smooth (“A-like”) colonies, it may indicate a durable infection (necessitating antibiotic use), whereas an isolate with only rough (“B-like”) or intermediate (“C-like”) morphotypes may indicate a weaker infection that may be resolve on its own or require a shorter antibiotic course. There is precedent for using morphotype composition to inform diagnosis and management of bacterial disease, as caused by another important opportunistic pathogen: *Pseudomonas aeruginosa*. *P. aeruginosa* is a leading cause of nosocomial infections, especially in individuals who are immunocompromised, and produces mucoid and rugose small-colony variants (SCVs) that differ in virulence (66). The different physical properties of *P. aeruginosa* morphotypes correspond to their production of distinct exopolysaccharides in the biofilm extracellular polymeric substance (67), and the presence of SCVs in the cystic fibrosis lung correlates with antibiotic resistance, declines in lung function, and prolonged infection (68).

The AFB staining properties of Component B and Component C, which respectively have mutations in *PKS* and *MtrA*, are consistent with the literature on these proteins’ involvement in cell wall lipidation and structure. PKSs are cytosolic enzymes which assemble into large multimeric, modular structures that catalyze the formation of numerous large molecules (69). In mycobacteria, the products of these biosynthetic factories include cell-wall associated lipids with extensive branched-chain fatty acids that are subsequently exported across the cell membrane by MmpLs (70, 71). Mycolactone, which causes the tissue destruction associated with Buruli ulcer, is also a PKS product (72). We predict the PKSΔCT deletion in Component B results in failed production of one or more cell-wall associated lipids, which in turn confer its rough colony appearance and weaker AFB staining. PKSs work in concert with MmpLs to then export these complex lipids into the extracellular space (73), which is presumably why Component B colonies resemble those of MmpL mutant *M. abscessus* (34, 35). Although our genome sequencing of *M. intracellulare ABSURDO* revealed it has fewer MmpL genes than anticipated, none of the morphotypes in our study had MmpL mutations, thus distinguishing ABSURDO components from the rough variants of *M. abscessus* (34, 35).

MtrA and MtrB make up a two-component system that is expressed in members of the phylum *Actinobacteria* (74). With MtrB being the sensor, MtrA is the response regulator which in turn activates downstream genes (75, 76). As such and in the case of ABSURDO, MtrA likely is not directly modifying the *M. intracellulare* cell wall but is instead promoting expression of proteins that are direct modifiers. Consistent with this, in *M. tuberculosis* the MtrA regulon includes a peptidoglycan remodeling enzyme, resuscitation-promoting factor, named so because it promotes the reactivation of dormant bacilli (77, 78). The absence of MtrA also impairs expression of cell wall homeostasis in *Dietzia*, another member of the phylum *Actinobacteria* which—like ABSURDO Component C—forms branched structures and is slow growing when MtrA is absent (79). In *M. intracellulare*, the genes which are downstream of MtrA have not been determined but would presumably be revealed by a future transcriptome comparison of COMPONENT C (in which MtrA is mutated) and COMPONENT A (in which MtrA is intact).

Regarding the point in time when these morphotypes arose we envision two possibilities: In the first, prior or at the time of initial infection, when the patient was first exposed to ABSURDO, each Component was already present in the infectious substance; since the etiology of NTM infection varies on a case-by-case basis we do not know if this infectious substance was soil, water or possibly even gastroesophageal reflux. As an alternative possibility, Component A was the only morphotype present at the infection event and subsequently gave rise to Component B and Component C in the patient lymph node. The genetic differences between *M. intracellulare* ABSURDO morphotype Components A-C provide an explanation for why both Component B and Component C are attenuated *in vitro* and *in vivo*, as disruptions to either lipid production (which may be expected in the absence of PKS) or peptidoglycan remodeling (which may occur in the absence of MtrA) would weaken the *M. intracellulare* cell wall. Since Component A grows the fastest, we propose its PKS and MtrA sequences be considered “wild type”, and the Component B-specific PKS sequence (ΔCT) and Component C-specific MtrA sequence (C→T) to be deleterious mutations. It should be noted, however, that we cannot rule out the contribution of other mutations in Component B or Component C which are predicted to have a Moderate effect on their encoded proteins (supplemental **TABLE S1** and **TABLE S2**), as testing this would require genetic complementation experiments. Molecular technologies for deleting genes in *M. intracellulare* have only recently been reported (80).

## ACKNOWLEDGEMENTS

We would like to acknowledge Dr. Susan Kehl of the Children’s Hospital of Wisconsin for providing the original isolate of *Mycobacterium intracellulare* ABSURDO. This work was supported in part by the Medical College of Wisconsin Research Affairs Committee Award (to A.R.H.). We would also like to acknowledge funding and support from The Ohio State University (to R.T.R and N.M.H.) and Bard College (to B.A.J.).

## SUPPLEMENTAL FIGURE LEGENDS

**FIGURE S1.**
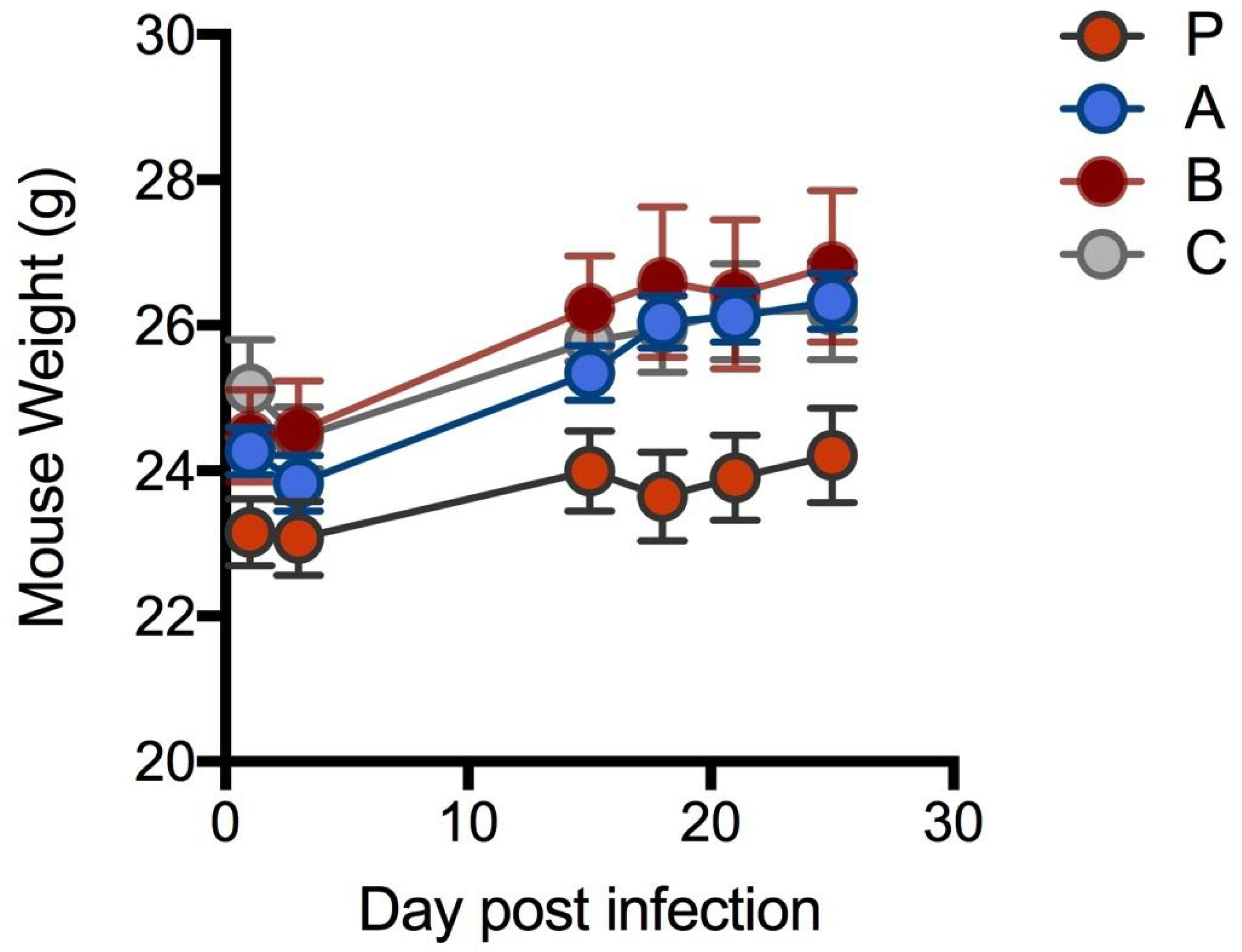
Weight change data of infected animals. The weight changes experienced by each group of B6 mice following inhalation exposure to either the parent isolate (P) of *M. intracellulare* ABSURDO or its morphotype components A, B or C.

**FIGURE S2.**
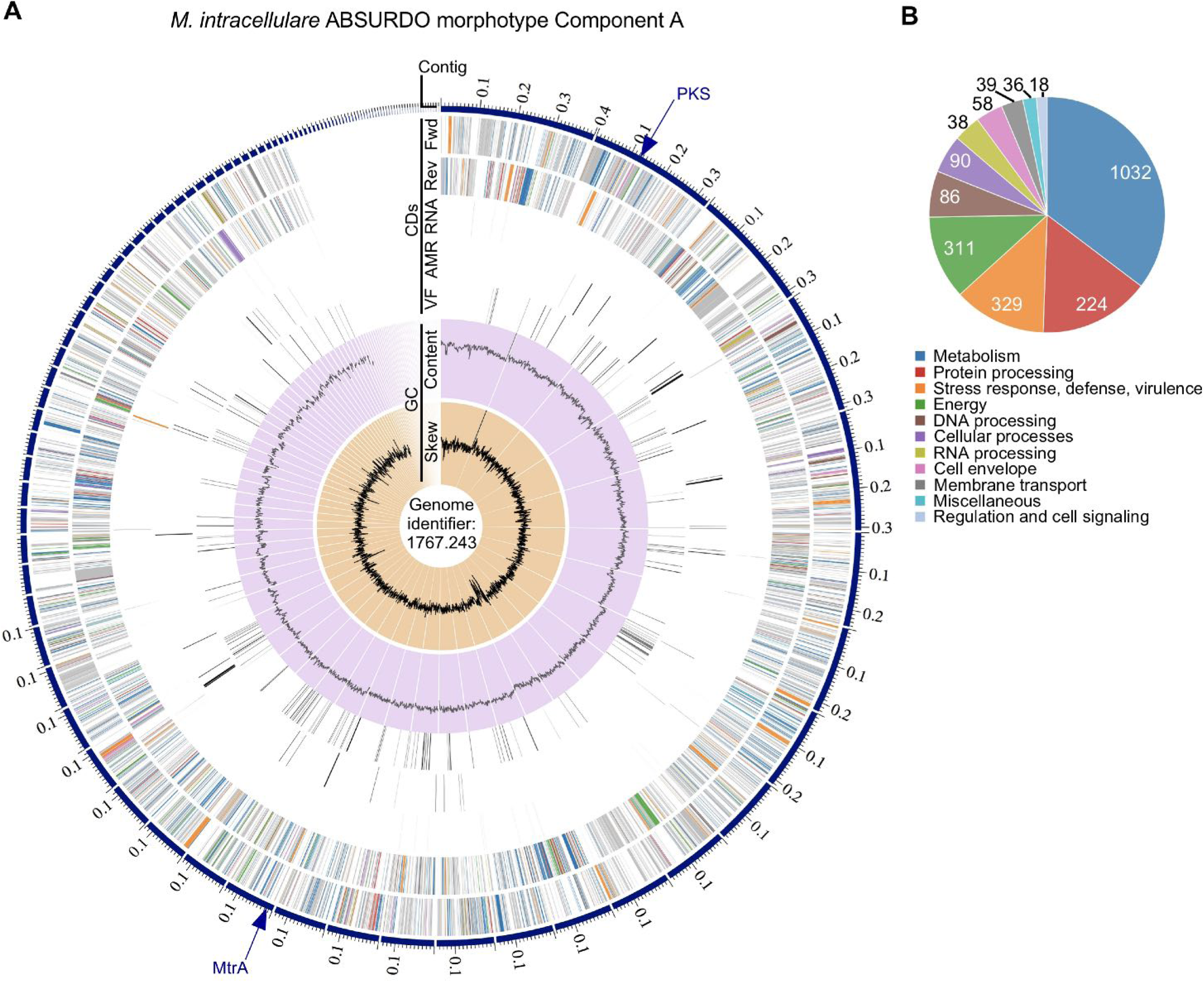
Annotated genome of the *M. intracellulare* ABSURDO morphotype Component A. Reads for *M. intracellulare* ABSURDO morphotype Component A were submitted to the comprehensive genome analysis service at PATRIC, analyzed using RAST tool kit, and—in the same manner as other genomes reported in this study—assigned a unique genome identifier of 1767.243. (**A**) A circular graphical display of the distribution of the genome annotations, which includes, from outer to inner rings, the contigs, CDS on the forward strand, CDS on the reverse strand, RNA genes, CDS with homology to known antimicrobial resistance genes, CDS with homology to know virulence factors, GC content and GC skew. The colors of the CDS on the forward and reverse strand correspond to the subsystem to which each gene belongs to per (**B**). Arrows indicate the locations of mutations in two genes—modular polyketide synthase PKS (location: contig 1767.243.con.0002 position 125064), and the two-component system response regulator MtrA (location: contig 1767.243.con.0018 position 10400)—that are the focus of FIG 7 and FIG 8, respectively, that distinguish Component A from the morphotypes Component B and Component C. (**B**) An overview of the subsystems for this genome, a subsystem being a set of proteins that together implement a specific biological process or structural complex. Values indicate the number of genes in the specified subsystem.

**FIGURE S3.**
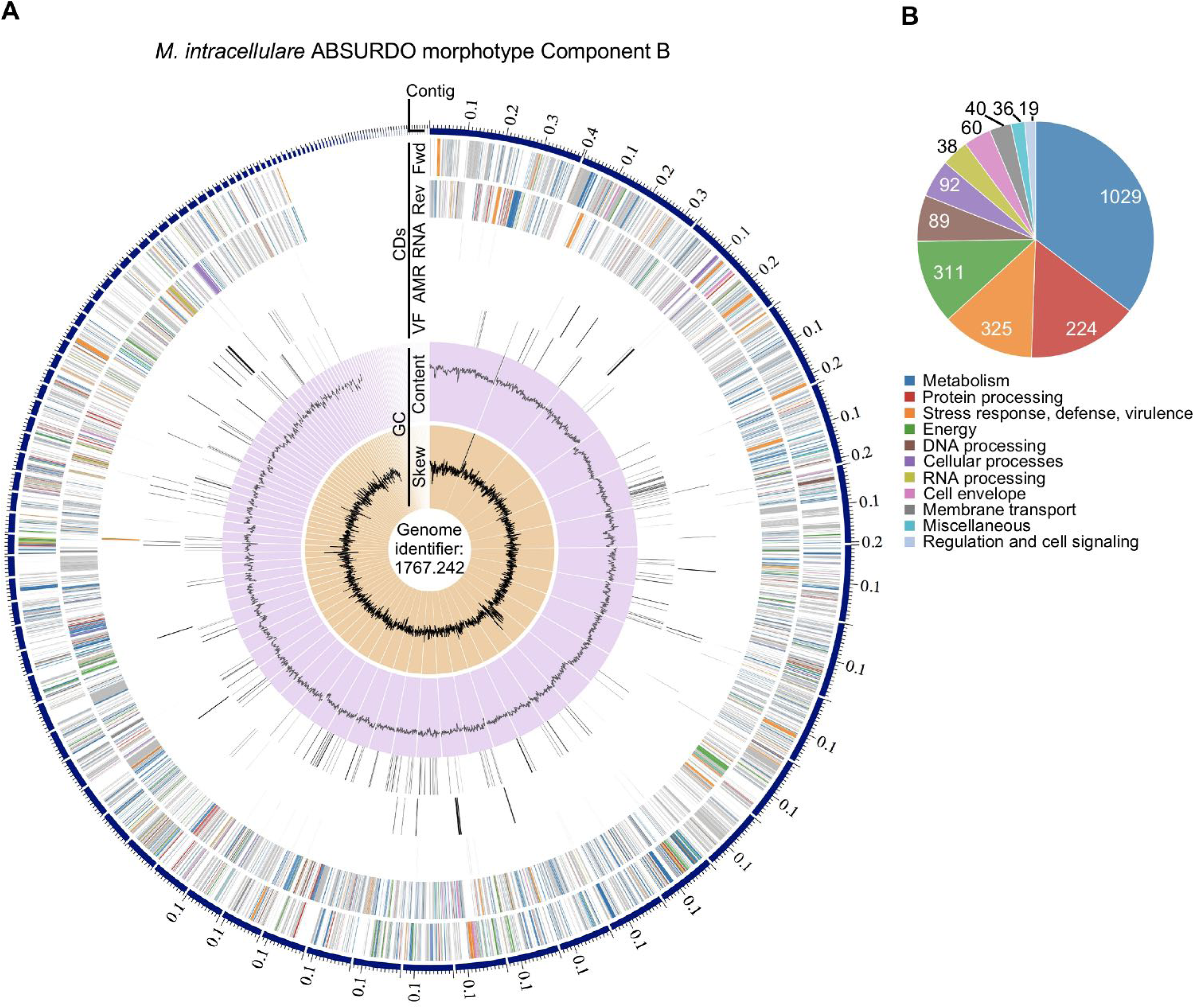
Annotated genome of the *M. intracellulare* ABSURDO morphotype Component B. Reads for *M. intracellulare* ABSURDO morphotype Component B were submitted to the comprehensive genome analysis service at PATRIC, analyzed using RAST tool kit, and—in the same manner as other genomes reported in this study—assigned a unique genome identifier of 1767.242. (**A**) A circular graphical display of the distribution of the genome annotations, which includes, from outer to inner rings, the contigs, CDS on the forward strand, CDS on the reverse strand, RNA genes, CDS with homology to known antimicrobial resistance genes, CDS with homology to know virulence factors, GC content and GC skew. The colors of the CDS on the forward and reverse strand correspond to the subsystem to which each gene belongs. (**B**) An overview of the subsystems for this genome, a subsystem being a set of proteins that together implement a specific biological process or structural complex. Values indicate the number of genes in the specified subsystem.

**FIGURE S4.**
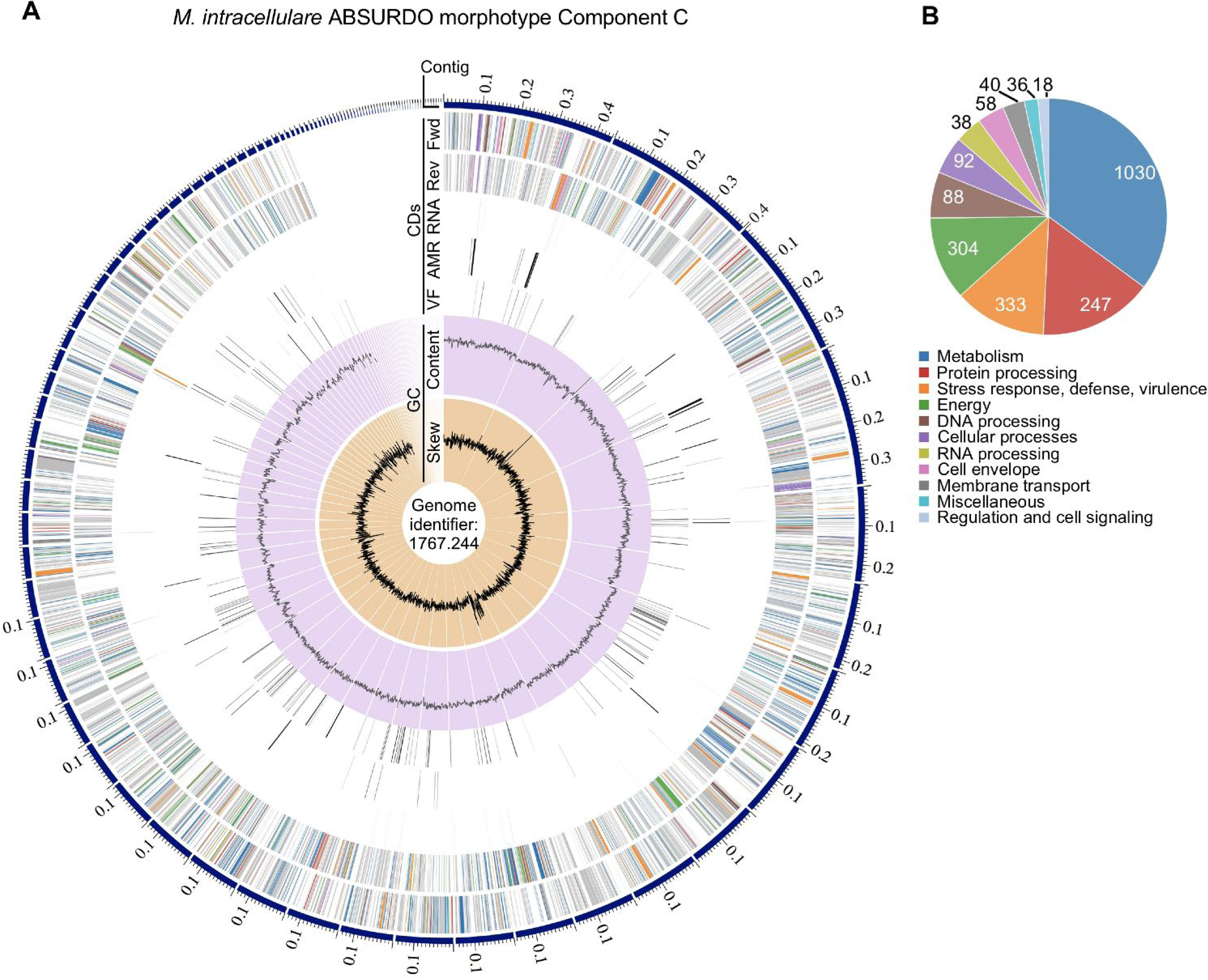
Annotated genome of the *M. intracellulare* ABSURDO morphotype Component C. Reads for *M. intracellulare* ABSURDO morphotype Component C were submitted to the comprehensive genome analysis service at PATRIC, analyzed using RAST tool kit, and—in the same manner as other genomes reported in this study—assigned a unique genome identifier of 767.244. (**A**) A circular graphical display of the distribution of the genome annotations, which includes, from outer to inner rings, the contigs, CDS on the forward strand, CDS on the reverse strand, RNA genes, CDS with homology to known antimicrobial resistance genes, CDS with homology to know virulence factors, GC content and GC skew. The colors of the CDS on the forward and reverse strand correspond to the subsystem to which each gene belongs. (**B**) An overview of the subsystems for this genome, a subsystem being a set of proteins that together implement a specific biological process or structural complex. Values indicate the number of genes in the specified subsystem.

**FIGURE S5.**
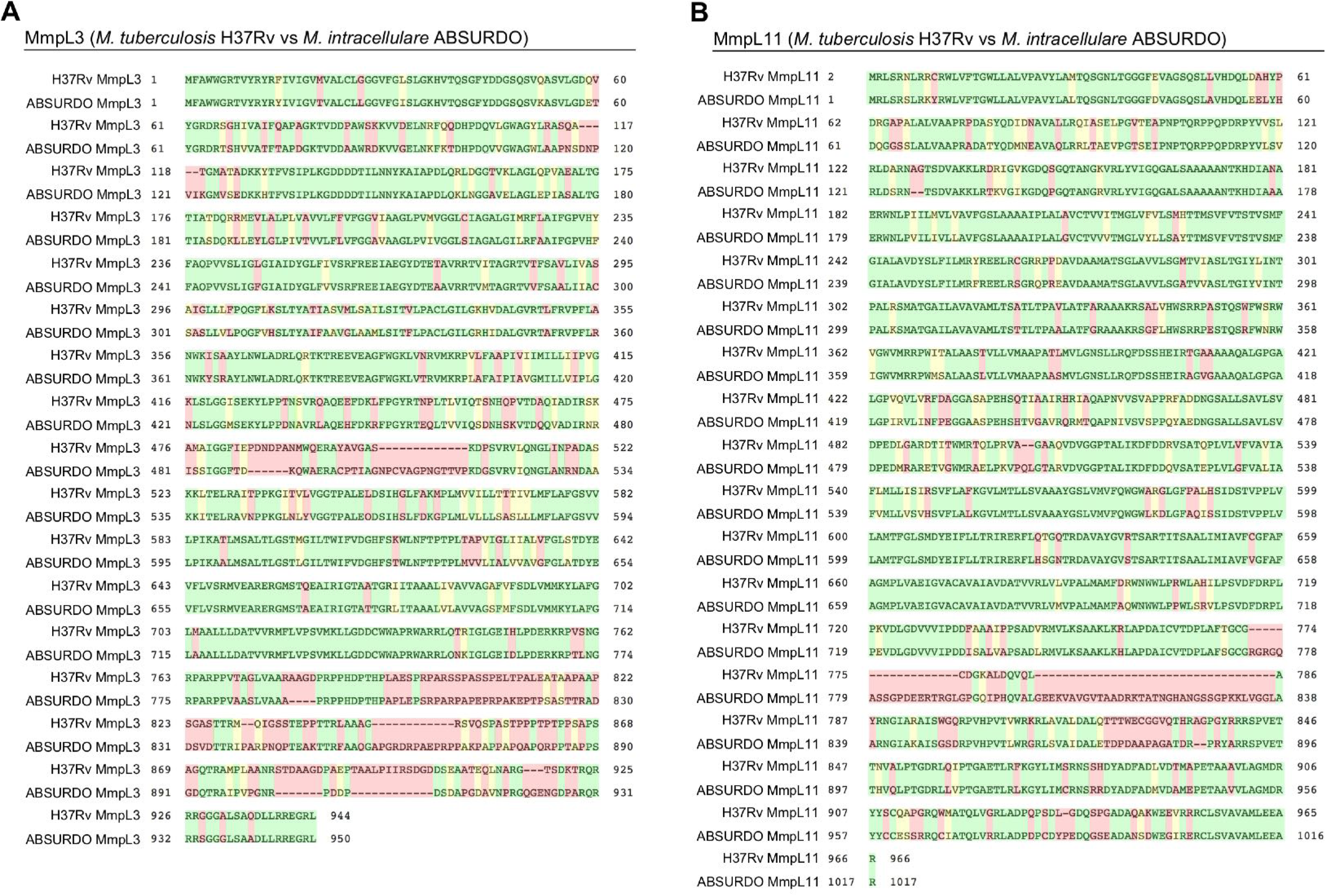
The *M. intracellulare* ABSURDO genome contains two genes that are homologous to *M. tuberculosis* MmpL3 and MmpL11. Two genes in the *M. intracellulare* ABSURDO genome, (A) *MmpL3* and (B) *MmpL11,* and the respective alignment to their *M. tuberculosis* H37Rv counterparts.

**FIGURE S6.**
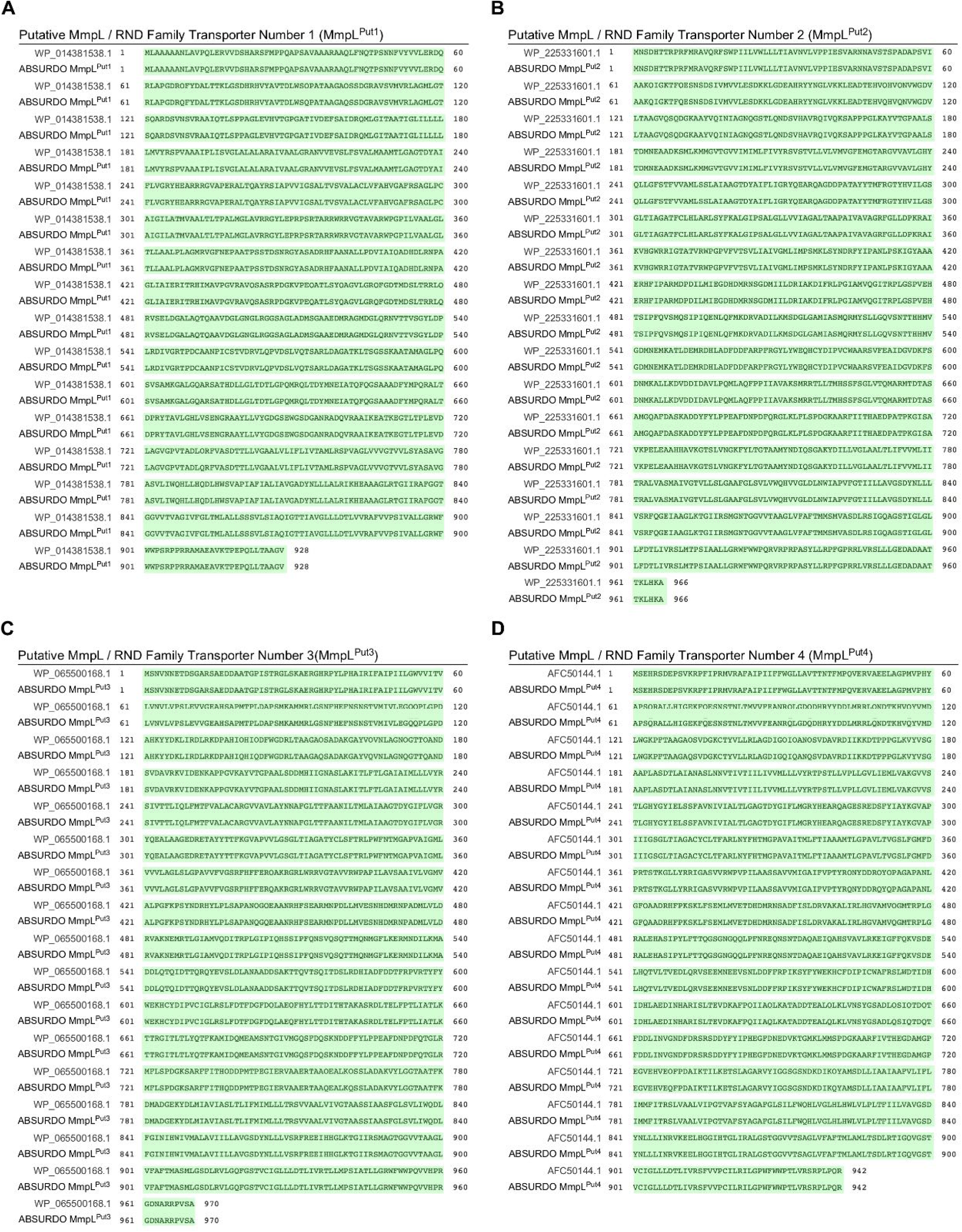
The *M. intracellulare* ABSURDO genome contains four additional genes that are putative MmpL / RND family transporters. Four putative MmpL/RND family transporter genes in the *M. intracellulare* ABSURDO genome, (A) *MmpL*^Put1^, (B) *MmpL*^Put2^, (C) *MmpL*^Put3^, and (D) *MmpL*^Put4^, and the respective alignment to the *M. intracellulare* NCBI gene IDs WP_014381538.1, WP_225331601.1, WP_065500168.1, and AFC50144.1.

**TABLE S1. *M. intracellulare* ABSURDO Component A Genome (1767.243) positions which differ from those in the Component B Genome (1767.242).** Listed from left to right are the genomic identifiers of each nucleotide difference between the genomes of Component A (reference genome) and Component B (variant genome), including the contig number, position, reference sequence, and variant sequence, as well as the predicted protein sequence changes associated with the Component C mutation (i.e. the ORF affected, its predicted function, the amino acid change, and whether the change has a low, moderate or high probability of impacting protein function). Relative to Component A, Component C has 18 ORFs affected by mutations, the majority of which were silent. Of those with non-synonymous mutations, only one resulted in a changed that was predicted to have a high impact on the encoding protein function: a frameshift mutation in modular PKS gene (the details of which are depicted in **FIG 7**).

**TABLE S2. *M. intracellulare* ABSURDO Component A Genome (1767.243) positions which differ from those in the Component C Genome (1767.244).** Listed from left to right are the genomic identifiers of each nucleotide difference between the genomes of Component A (reference genome) and Component C (variant genome), including the contig number, position, reference sequence, and variant sequence, as well as the predicted protein sequence changes associated with the Component C mutation (i.e. the ORF affected, its predicted function, the amino acid change, and whether the change has a low, moderate or high probability of impacting protein function). Relative to Component A, Component C has 16 ORFs affected by mutations, the majority of which were silent. Of those with non-synonymous mutations, only one resulted in a changed that was predicted to have a high impact on the encoding protein function: a stop codon in the two-component system response regulator MtrA (the details of which are depicted in **FIG 8**).

